# The activities of LRRK2 and GCase are positively correlated in clinical biospecimens and experimental models of Parkinson’s disease

**DOI:** 10.1101/2021.09.27.461935

**Authors:** Maria Kedariti, Emanuele Frattini, Pascale Baden, Susanna Cogo, Laura Civiero, Elena Ziviani, Massimo Aureli, Alice Kaganovich, Mark R Cookson, Leonidas Stefanis, Matthew Surface, Michela Deleidi, Alessio Di Fonzo, Roy N. Alcalay, Hardy Rideout, Elisa Greggio, Nicoletta Plotegher

## Abstract

LRRK2 is a kinase involved in different cellular functions, including autophagy, endolysosomal pathways and vesicle trafficking. Mutations in *LRRK2* cause autosomal dominant forms of Parkinson’s disease (PD). Heterozygous mutations in *GBA1*, the gene encoding the lysosomal enzyme glucocerebrosidase (GCase), are the most common genetic risk factors for PD. Moreover, GCase function is altered in idiopathic PD and in other genetic forms of the disease. Recent work suggests that LRRK2 kinase activity can regulate GCase function. However, both a positive and a negative correlation have been described. To gain insights into the impact of LRRK2 on GCase, we investigated GCase levels and activity in *LRRK2* G2019S knockin mice, in clinical biospecimens from PD patients carrying this mutation and in patient-derived cellular models. In these models we found a positive correlation between the activities of LRRK2 and GCase, which was further confirmed in cell lines with genetic and pharmacological manipulation of LRRK2 kinase activity. Overall, our study indicates that LRRK2 kinase activity affects both the levels and the catalytic activity of GCase.

## Introduction

Mutations in *LRRK2* cause autosomal dominant Parkinson’s disease (PD) with age- and mutation-dependent penetrance^1–3^, whereas heterozygous mutations in *GBA1* are the most common genetic risk factors for PD and the cause of the lysosomal storage disorder Gaucher disease when present in homozygosis^4,5^. LRRK2 is a large, multi-domain protein with two enzymatic domains, a Ser/Thr kinase domain and a small GTPase domain (ROC), where the bulk of the pathogenic PD-linked mutations are located. While its full range of cellular functions has yet to be characterized, it has been robustly associated with endo-lysosomal pathways and vesicular trafficking (reviewed in Bonet-Ponce and Cookson, 2021^6^). These activities are likely mediated by its phosphorylation of multiple members of the Rab GTPase family, which is increased in the context of the disease-linked mutations^7^, and potentially also in cases of PD not linked to mutations in *LRRK2*^8^.

The main function of the lysosomal enzyme glucocerebrosidase (GCase) is to hydrolyse glucosylceramide and glucosylsphingosine to glucose and either ceramide or sphingosine respectively; and most of the mutations in *GBA1* associated with PD risk reduce the activity of GCase^5,9^. High levels of α-synuclein, another protein mutated in PD and the major component of Lewy bodies, inhibit autophagic flux and the lysosomal activity of GCase^10^. GCase activity has been shown to be reduced also in peripheral monocyte extracts from PD patients without mutations in *GBA1*^11,12^ and in PD brains^13^, overall suggesting that alterations of GCase activity may be a common underlying feature of PD, similar to what has been proposed for changes in LRRK2 kinase activity.

Using a novel method of assessing GCase activity in dried blood spots, in 2015 Alcalay and colleagues reported a significant increase in GCase activity in carriers of the *LRRK2* G2019S mutation^14^, suggesting that carriers of the gain of function mutation have higher GCase activity and therefore the activities of the two enzymes are positively correlated in blood cells. Shortly after, using brain lysates from LRRK2 knock-out (KO) mice, we found that loss of LRRK2 results in decreased GCase levels, which corresponded to an increase in GCase specific activity^15^. Because of the role played by LRRK2 in the vesicular and endo-lysosomal systems, several studies have followed to assess the link between mutant LRRK2 and GCase activities.

Recently, GCase activity of induced pluripotent stem cell (iPSC)-derived dopamine neurons from LRRK2-PD patients, carrying either the G2019S or the R1441C mutations, was found to be reduced compared to neurons from healthy controls, suggesting a negative correlation between these two activities^16^. There may be several possibilities for the discrepancy between this result and the results obtained from blood samples. First, there may be a cell-type specific link between the activities of LRRK2 and GCase, manifesting in divergent ways in neuronal cells compared to peripheral blood immune cells. Second, methodological differences in the assessment of GCase activity (i.e., cell based vs. *in vitro*) may capture different aspects of this interplay. A further study on blood samples from PD-patients identified a correlation between the *LRRK2* variant M1646T and increased GCase activity^17^, consistent with the result we had obtained for *LRRK2* G2019S carriers^14^.

Interestingly, the interplay between LRRK2 and GCase was also identified in iPSC-derived astrocytes derived from PD patients carrying *GBA1* mutations, in which LRRK2 inhibition could rescue the lysosomal and inflammatory defects^18^.

To shed more light into the relationship between *LRRK2* and *GBA1*, here we assessed GCase activities in multiple LRRK2 cellular and *in vivo* models, including plasma and PBMC samples from idiopathic PD and *LRRK2* G2019S cohorts. Our findings support a model in which increased LRRK2 kinase activity leads to enhanced intrinsic GCase activity.

## Results

### G2019S LRRK2 mutation impacts GCase function in mouse brain

To understand the effect of LRRK2 mutations on GCase function in brain tissues, we performed GCase activity assays and western blot analysis on brain lysates from different regions (midbrain, striatum and cortex) in 6-month-old G2019S knockin (GSKI) and WT mice. GCase activity normalized by total proteins is different across brain regions, but not across genotypes (Fig. 1a, two-way ANOVA, Bonferroni post-test; brain region, P = 0.0015; genotype, P = 0.0708; interaction, P = 0.7089; n = 4-5). Specifically, the GCase activity measured in lysates from midbrain and cortex is higher compared to that observed in the striatum (Fig. 1a). Considering that dopaminergic neurons in the midbrain are the most vulnerable in PD, we focused on this region to further evaluate GCase levels. Strikingly, GCase protein levels are significantly lower in GSKI midbrain lysates as compared to WT (Fig. 1b). The specific GCase activity (i.e., activity normalized by the amount of enzyme) in GSKI midbrains was significantly higher than in WT midbrains (Fig. 1c).

**Figure 1.**
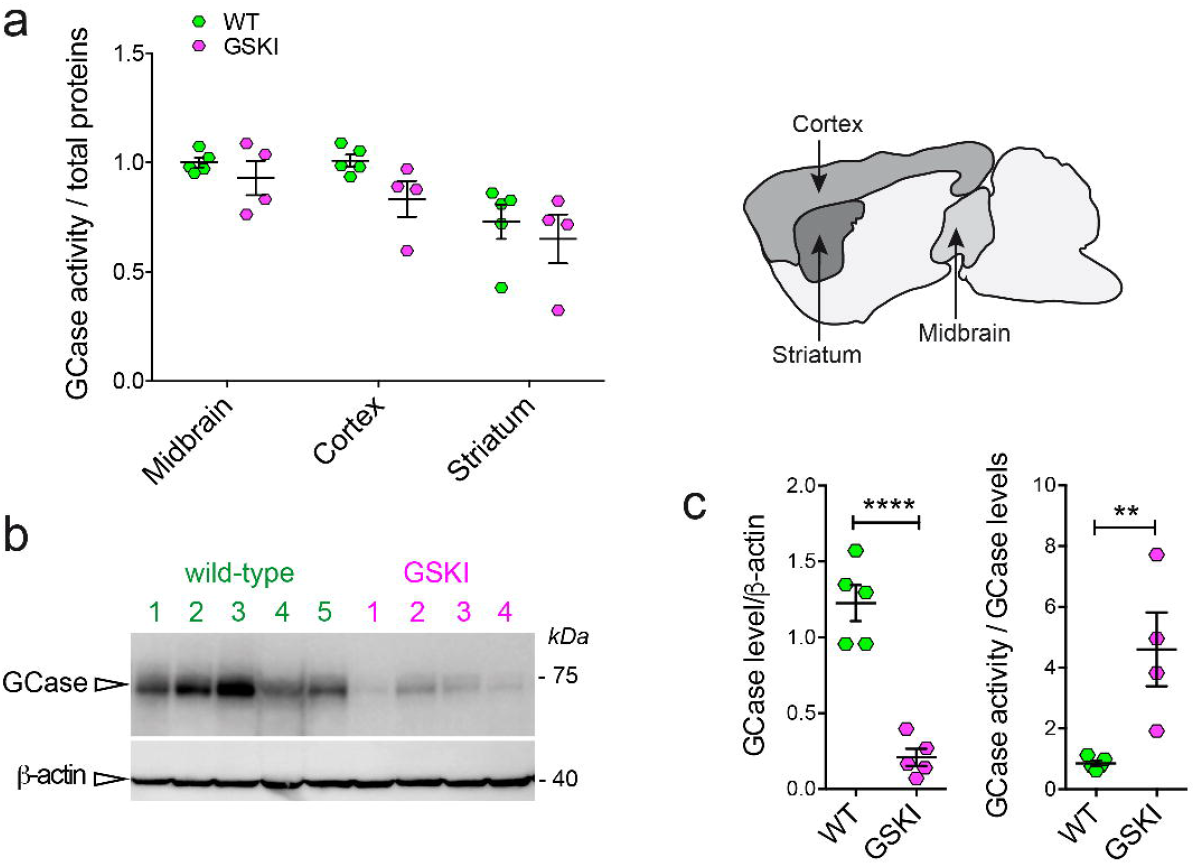
GCase activity and level measured in *LRRK2* G2019S KI mice brains. a. GCase activity measured in brain extracts from different brain regions (midbrain, cortex and striatum) shows limited differences between WT and GSKI brains, while the difference is significant across the different measured tissue extract (n=4-5 samples per genotype, two-way ANOVA test, p=0.015 for brain regions and P=0.0708 for genotype). b. Western blot analysis of GCase protein levels and the relative β-actin loading control of tissue extracts from the midbrain for WT and KI mice. c. Quantification of the GCase level normalized to β-actin showed a significant decrease in GSKI midbrain compared with WT tissue extracts, while GCase activity was significantly higher in GSKI mice tissue when normalized to the GCase level (n=4-5 samples per genotype, Student t test, **** p<0.00001; ** p<0.001).

### GCase activity is increased in affected LRRK2 G2019S mutation carriers

It has been previously reported that GCase activity is elevated in *LRRK2* G2019S carriers using a novel dried blood spot assay^14^. Here we tested if these results can be replicated in purified PBMCs from a similar cohort, hypothesizing that GCase measurements from PBMCs would be more accurate than measurements in dried blood spots. In this cohort, we previously found that *in vitro* LRRK2 kinase activity purified from PBMCs is significantly increased in both affected and healthy carriers of the *LRRK2* G2019S mutation compared to healthy controls and PBMCs from idiopathic PD (iPD) patients^19^.

As expected, GCase activity in PBMCs from carriers of *GBA1* mutations was significantly lower compared to PBMCs from healthy control subjects (Fig. 2a). Likewise, as reported previously^20^, the GCase activity in PBMCs from iPD patients was not significantly different from healthy controls (Fig. 2a). Interestingly, we found that GCase activity is increased in PBMCs from G2019S carriers, in agreement with our previous study^14^ (Fig. 2a). This was evident only in PBMCs from affected carriers of the *LRRK2* G2019S mutation, as non-manifesting carriers exhibit GCase activity similar to controls. We had insufficient amounts of PBMC extracts normally required for the evaluation of GCase protein level and for crude isolation of lysosomes; thus, for these experiments, the fluorescent signal arising from GCase activity in the whole lysate was normalized to total protein expression. It is worth noting that the distribution of GCase activities in symptomatic G2019S carriers is larger compared to the other groups. This could be due to sampling variance, but could also suggest that when PD manifests, the G2019S mutation increases the likelihood that GCase activity is enhanced. Thus, in this mixed immune cell population, the link between LRRK2 kinase activity and GCase activity is not strictly tied to either LRRK2 mutation status, or disease status alone, but rather a synergistic effect between the two may be needed to alter the intrinsic activity of GCase.

**Figure 2.**
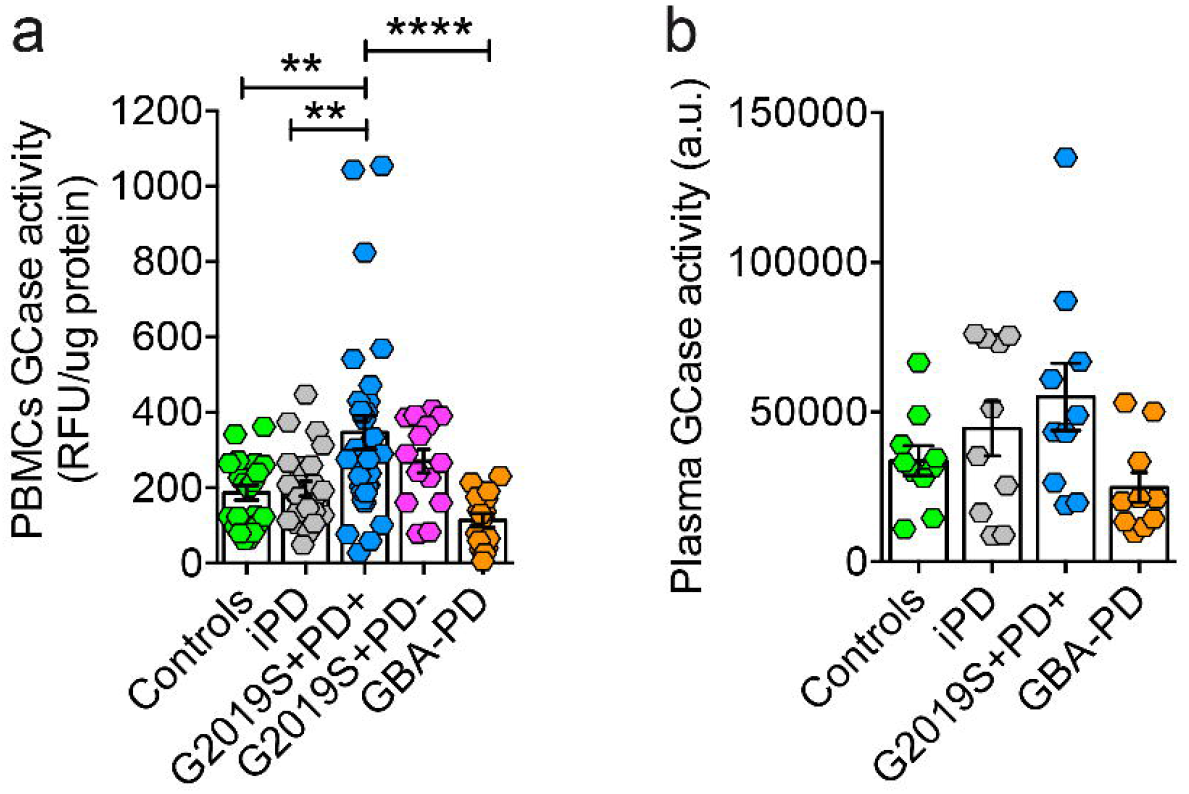
GCase activity and level measured in PBMCs and plasma from idiopathic PD patients, in *LRRK2* G2019S unaffected carriers and patients, in *GBA1* PD patients compared with controls. a. GCase activity (normalized to total protein content) measured in PBMC extracts from the indicated PD cohort. While GCase activity in GBA1 mutation carriers is reduced, activity in affected G2019S-LRRK2 carriers is significantly elevated. (One-way ANOVA, Tukey’s post-hoc tests were performed; ** p<0.01, **** p<0.0001). b. GCase activity normalized by the total protein content measured in the plasma of idiopathic patients, in patients carrying *LRRK2* G2019S mutations or *GBA1* mutations. The trend (although not statistically significant, one-way ANOVA, Tukey’s post-test) is similar between plasma samples and PBMCs (10 individuals for each group).

A similar trend was obtained by measuring GCase activity in plasma from a small pilot cohort (see Material and Methods) constituted by *LRRK2* G2019S PD patients, *GBA1* PD patients, iPD patients and controls (Fig. 2b).

### Endolysosomal compartment defects correlate with GCase activity in LRRK2 G2019S patient fibroblasts

To further investigate the role of the hyperactive *LRRK2* G2019S mutant on GCase behavior in another type of patient-derived cells, we analyzed PD-patient fibroblasts carrying G2019S LRRK2 mutation. Three patient cell lines were first analyzed by Transmission Electron Microscopy (TEM) in comparison with age and sex matched controls, to identify macroscopic alterations in the endo-lysosomal compartment in relationship to LRRK2 mutation and GCase activity. Representative TEM micrographs reported in Fig. 3a show a striking accumulation of electron dense or lamellar structures in LRRK2 patient cells. They are likely endo-lysosomes engulfed with undigested materials, autophagic vacuoles and multilamellar bodies, which are all typical hallmarks of altered autophagic lysosomal pathways. Interestingly, multilamellar bodies (MLBs) were also found in fibroblasts from PD-patients carrying the N370S mutation in the GBA gene^21,22^. Quantification of the number of MLBs per area across the different cell lines revealed an increase in patient lines compared to controls, but still G2019S carrier lines showed a higher degree of variability (Fig. 3b-c). These morphological alterations are not accompanied by an altered level of the lysosomal marker LAMP-1 as evaluated by western blot analysis, nor by a significant change in the activity or level of GCase in the patient lines compared to controls, likely because of the high variability among individuals (Fig. 3d-e). Nevertheless, in *LRRK2* G2019S fibroblasts, the trend in terms of GCase activity and level is similar to the one found in the GSKI brains and in G2019S blood samples from PD patients. Moreover, there is a trend suggesting a negative correlation between GCase activity/GCase protein level and the amount of MLBs present in patient fibroblasts, while the same does not occur for controls (Fig. 3f).

**Figure 3.**
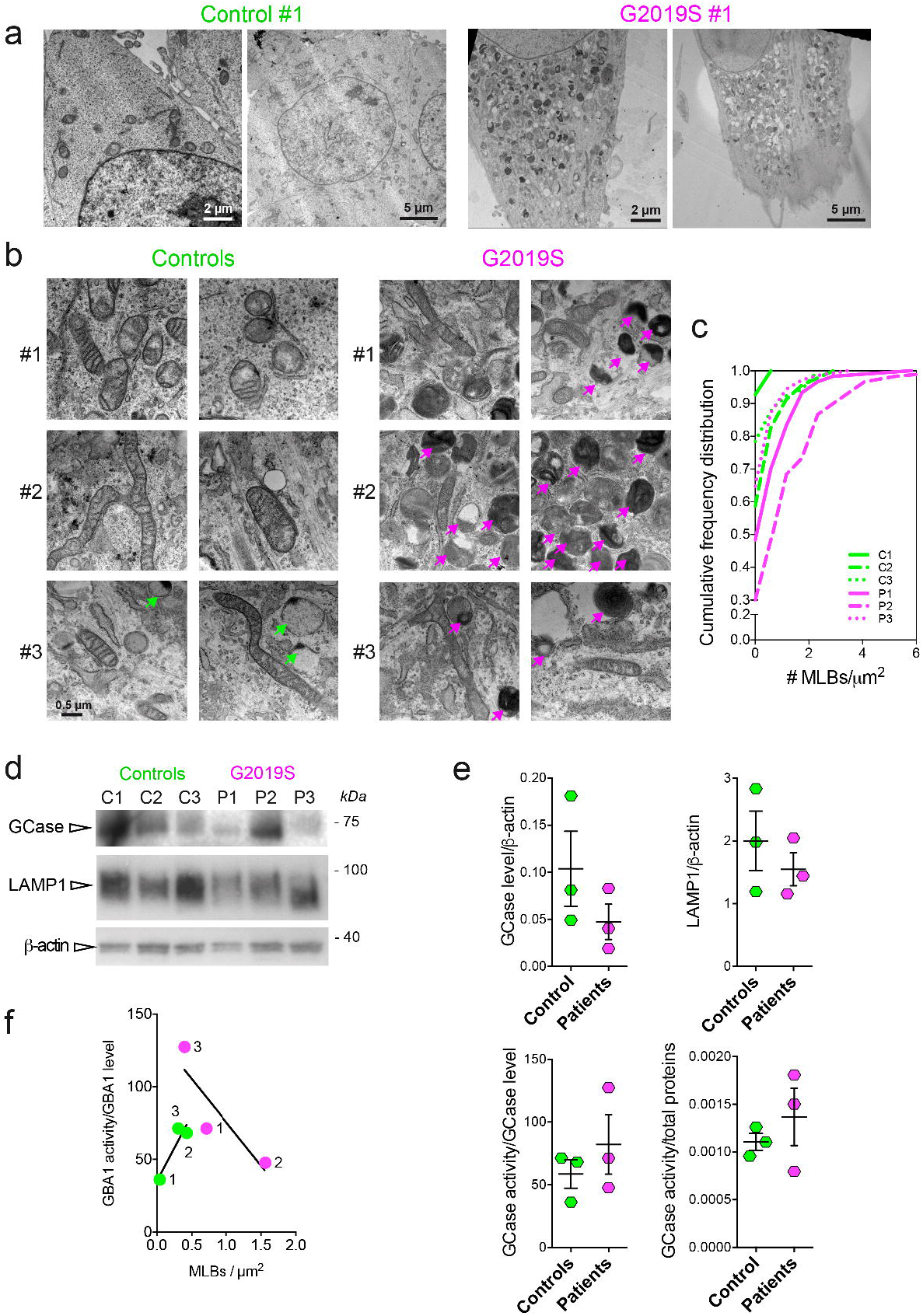
Ultrastructural analysis of endo-lysosomal compartment and GCase function in fibroblasts from *LRRK2* G2019S mutant compared with controls. a. Low magnification (2-5 µm) TEM micrographs showing electron-dense structures accumulating in the *LRRK2* G2019S fibroblasts (Patient #1), while they are almost completely absent in control cells (Control #1). b. High magnification (500 nm) TEM images three patient fibroblast cell lines and age and sex matched controls. Patient fibroblasts show accumulation of endo-lysosomal structures that resemble multilamellar bodies (MLBs). c. Cumulative distribution of the number of the MLBs per µm^2^ for each patient and each control cell line, showing that the number of MLBs is larger in *LRRK2* G2019S fibroblasts than in control cells (given a certain degree of variability across individuals). d. Western blot analysis (each dot represents the average of n=3 technical replicates) of GCase and LAMP1 levels in three *LRRK2* G2019S fibroblast lines and in three related controls. e. Quantification of the GCase and LAMP1 protein levels as measured by western blot and normalized by β-actin; GCase activity normalized by total protein or by GCase protein levels. This measurement shows no significant difference between LRRK2-PD patients and controls. f. Negative correlation between the intrinsic GCase activity (i.e., GCase activity/GCase protein level) and the number of MLBs is observed in *LRRK2* G2019S patients while is absent in control fibroblast lines (r = 0.92 for control cells and r = -0.88 for patients’ cells, Pearson correlation analysis).

### GCase activity is increased in iPSC-derived dopaminergic neurons from G2019S LRRK2 patients

We next evaluated the role of LRRK2 kinase activity on GCase function in a human-derived disease-relevant model. To this aim, we took advantage of iPSC-derived dopaminergic neurons. iPSC lines of PD patients and controls stained positive for stem cell markers (SOX2, OCT4, TRA-1-60, SSEA4). Karyotype analysis excluded genetic abnormalities of reprogrammed iPSCs. Dopaminergic neurons differentiated from all iPSC lines cultured up to 70 days in vitro stained positive for neuronal (TUJ1) and dopaminergic (TH, DAT) markers, confirming appropriate differentiation (Fig. 4a).

**Figure 4.**
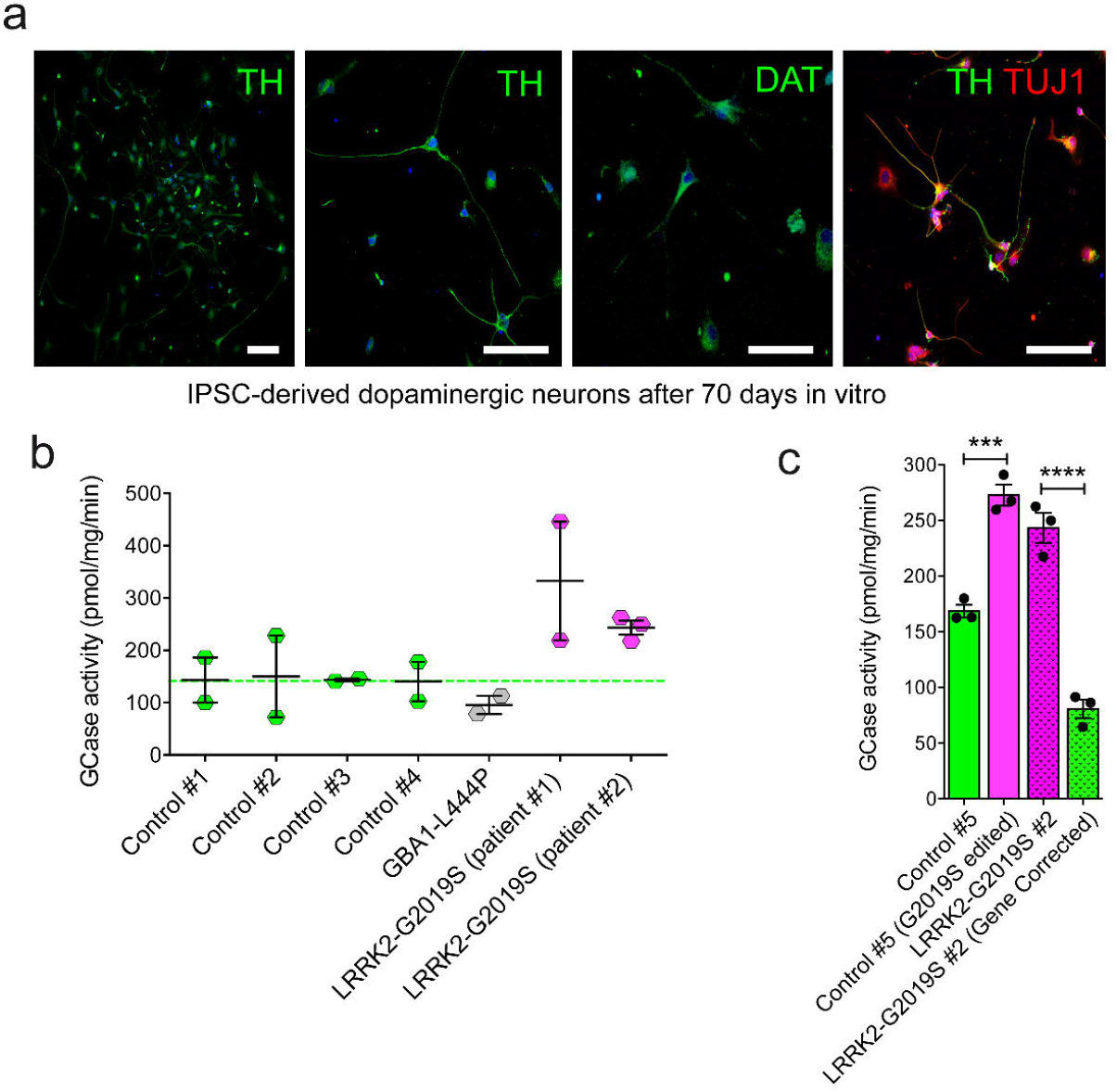
iPSc-derived neurons from *LRRK2* G2019S PD patients and *GBA1* PD patient show altered GCase activity compared to neurons derived from healthy subjects. a. Representative confocal images of iPSC-derived neurons stained with TH, DAT and TUJ1 antibodies to verify proper differentiation to dopaminergic cells. b. GCase activity measured in iPSC-derived neurons obtained from four healthy controls, one *GBA1* L444P PD patients and two *LRRK2* G2019S PD patients (2-3 replicates per individual). Data show that *GBA1* PD patients have a reduced GCase activity compared with controls, while *LRRK2* PD patients present increased GCase activity. c. GCase activity measured in iPSC-derived dopaminergic neurons obtained from one healthy control in which the *LRRK2* G2019S was introduced (Control #5 G2019S edited) show an increased enzymatic activity compared with the non-edited neuronal cells (Control #5). Similarly, GCase activity was reduced to a level similar to the Control #5 iPSC-derived neurons in LRRK2 G2019S dopaminergic cells (LRRK2-G2019S #2) upon gene correction (LRRK2-G2019S #2 GC). One-way ANOVA with Tukey’s post-test (**p < 0.01; ****p < 0.0001).

As a first screen, we measured GCase activity in iPSC-derived neurons carrying mutations in different PD genes, i.e., *GBA1* (L444P), *LRRK2* (G2019S) and compared to 5 different healthy controls. In line with results obtained in PBMCs, and as expected, iPSC-derived dopaminergic neurons carrying L444P mutation in *GBA1* gene showed decreased activity of GCase, compared to healthy controls (Fig. 4b). Consistent with data on GCase activity in PBMCs from PD patients carrying G2019S *LRRK2* mutation, two lines of iPSC-derived dopaminergic neurons carrying the G2019S mutation in *LRRK2* gene showed increased GCase activity (Fig. 4b). To demonstrate that the effect on GCase activity is genuinely mediated by the *LRRK2* G2019S mutation, dopaminergic neurons were differentiated from an isogenic control line with wild type LRRK2 (LRRK2 #2 GC) corrected from iPSCs of a *LRRK2* G2019S patient (LRRK2 #2) and from an isogenic line from a control subject (CTR5) edited to carry *LRRK2* G2019S mutation (CTR5 G2019S). The increase in the GCase activity observed in the LRRK2 mutated line was significantly lowered in the isogenic control with corrected LRRK2. Similarly, the introduction of the *LRRK2* G2019S mutation in the control line resulted in an increased GCase activity (Fig. 4c). Taken together, we collected multiple lines of evidence that GCase activity is increased in different cell types, including iPSCs-derived dopaminergic neurons and peripheral immune cells, expressing hyperactive G2019S LRRK2.

### Genetic or pharmacological inhibition of G2019S LRRK2 in HEK293T cells leads to an increase in GCase levels, but a decrease in GCase activity

We next examined if the effect of the *LRRK2* G2019S mutation on GCase activity specifically depends upon the increased kinase activity associated with this mutation. To this aim, we employed HEK293T cells transiently over-expressing *LRRK2* G2019S, and evaluated GCase activity and level upon pharmacological inhibition of LRRK2 kinase activity using MLi-2. As shown in Fig. 5a, 100 nM MLi-2 inhibition for 90’ or 5 hours caused a reduction of GCase activity normalized by GCase level by 50% of the activity measured in untreated *LRRK2* G2019S over-expressing cells. This is due to a combination between the slight decrease in GCase activity when normalized by the total protein concentration (Fig. 5b) and the increase in GCase protein level (Fig. 5a, 5b) as measured by western blot analysis (n=4 independent experiments, One-way ANOVA test, Bonferroni post-hoc test). The GCase inhibitor conduritol-β-epoxide (CBE) was used as a positive control and efficiently reduced GCase activity upon 5 hours treatment, without significantly affecting the protein level and the effect of MLi-2 was confirmed by the reduction of LRRK2 phosphorylation at Ser935 (Fig. 5a).

**Figure 5.**
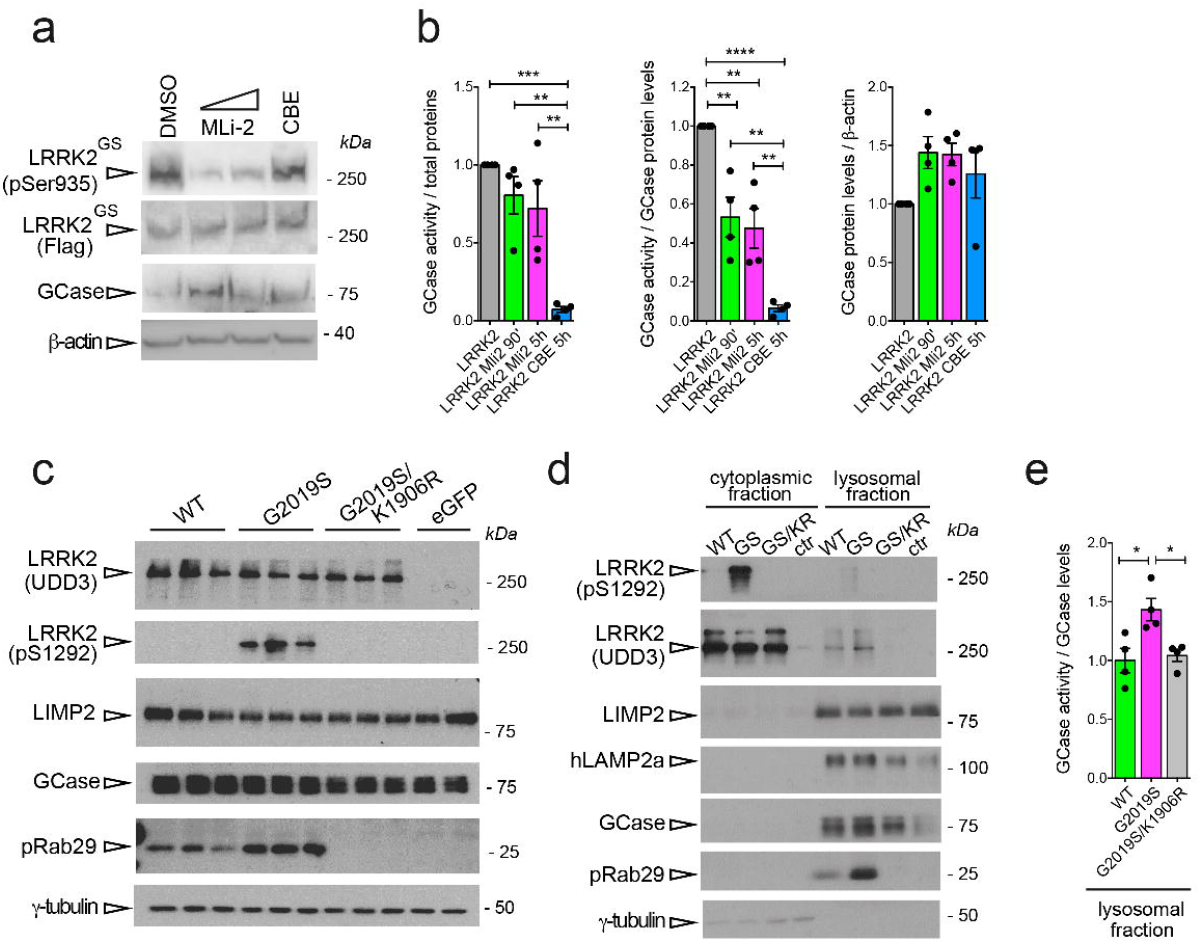
GCase activity and level is altered in HEK293T cells in which LRRK2 kinase activity is pharmacologically or genetically manipulated. a. Representative western blot of LRRK2-flag, pSer935 LRRK2 and GCase activity in HEK293 cells overexpressing LRRK2 G2019S and treated with the LRRK2 inhibitor Mli2 for 90’ or 5 hours, or with the GCase inhibitor CBE for 5 hours. b. Quantification of the GCase activity (normalized by total proteins or by GCase levels) show that the pharmacological inhibition of LRRK2 activity reduces GCase activity in both cases, even though to a lesser extent compared with CBE treatment. GCase protein level is slightly increased upon both MLi-2 and CBE treatment, though non-significantly. One-way ANOVA with Tukey’s post-test (**P < 0.01; ***P < 0.001; ****P < 0.0001). c. Representative western blot of HEK293T cells over-expressing WT, G2019S, or G2019S/K1906R (kinase-dead) LRRK2 probed for total (UDD3) and phosphorylated (pS1292-LRRK2) LRRK2, LIMP1, GCase, phosphorylated Rab29 (T71), and γ-tubulin. LRRK2 kinase function, increased autophosphrylation and phosphorylation of endogenous Rab29 in cells expressing G2019S-LRRK2 is prevented by the kinase inactivating mutation (K1906R). d. Representative western blot of cytoplasmic and lysosomal fraction of HEK293T cells over-expressing WT, G2019S, or G2019S/K1906R (kinase-dead) LRRK2 probed for total (UDD3) and phosphorylated (pS1292-LRRK2) LRRK2, LIMP1, GCase, phosphorylated Rab29 (T71), and γ-tubulin. e. Quantification of GCase activity normalized by GCase level in the lysosomal fraction of HEK293T cells over-expressing WT, G2019S, or G2019S/K1906R (kinase-dead) LRRK2 showed a significant increase in G20129S samples compared with LRRK2 WT or with the kinase-dead mutant. (One-way ANOVA with Tukey’s post-test; *p < 0.05).

As a complementary approach, we transiently overexpressed LRRK2 WT, mutant LRRK2 G2019S, or a LRRK2 kinase-dead double mutant G2019S/K1906R, in HEK293T cells and assessed GCase activity in crude lysosomal fractions. To confirm that the over-expressed G2019S and G2019S/K1906R mutants elicited the expected effects on LRRK2 kinase activity, we assessed auto-phosphorylation of LRRK2 at Ser1292. Cells expressing LRRK2 G2019S exhibit increased auto-phosphorylation at Ser1292, as expected, which is reversed by the kinase-dead form of this mutant (Fig. 5c).. In crude lysosomal fractions, we could detect total LRRK2 (Fig. 5d), as well as a faint band corresponding to pS1292-LRRK2 in lysosomes of cells expressing *LRRK2* G2019S, and increased pT71-RAB29. The lysosomal localization of both phosphorylated as well as total LRRK2 was dependent on its kinase activity, as the K1906R inactivating mutation blocked these accumulations (Fig. 5d).

To determine if LRRK2 and GCase activities are linked in this cellular model, we prepared a crude lysosomal fraction and assessed GCase activity as above. Similar to what we observed in other models, we found a significant increase in GCase activity in enriched lysosomal fractions of cells expressing *LRRK2* G2019S, as normalized to GCase levels (Fig. 5e). Again, the kinase-dependency of this induction was confirmed by the reversal of this phenotype in cells expressing kinase-dead G2019S/K1906R LRRK2.

### Lysosomal GCase activity in LRRK2 knockout macrophages

To further complement and expand these findings, we took advantage of the macrophage cell model RAW264.7 where LRRK2 biology is well-characterized ^23,24^. One advantage of this model is that macrophages express both GCase and LRRK2 at high levels, allowing to study the impact of LRRK2 kinase activity on GCase function without over-expressing LRRK2. Moreover, in this cell type, LRRK2 exerts crucial functions in the regulation of immune and inflammatory responses^24^, but also in the control of lysosomal damage^25,26^. Here, we assessed GCase activity using the 4-MU substrate based-assay in crude lysosomal preparations isolated from LRRK2 WT and knockout (KO) macrophages and compared it with GCase activity measured in cells using a live cell-based fluorescent assay (PFB-FDGlu)^11,16^. Both GCase levels and GCase activity assessed with the 4-MU substrate are reduced by 50% in crude lysosomal preparations from LRRK2 KO cells (Fig. 6a-b). The graph in Fig. 6c shows a decrease in GCase protein level in LRRK2 KO (∼ 60%) and a non-significant increase in GCase activity normalized to GCase levels in total cell extracts.

**Figure 6.**
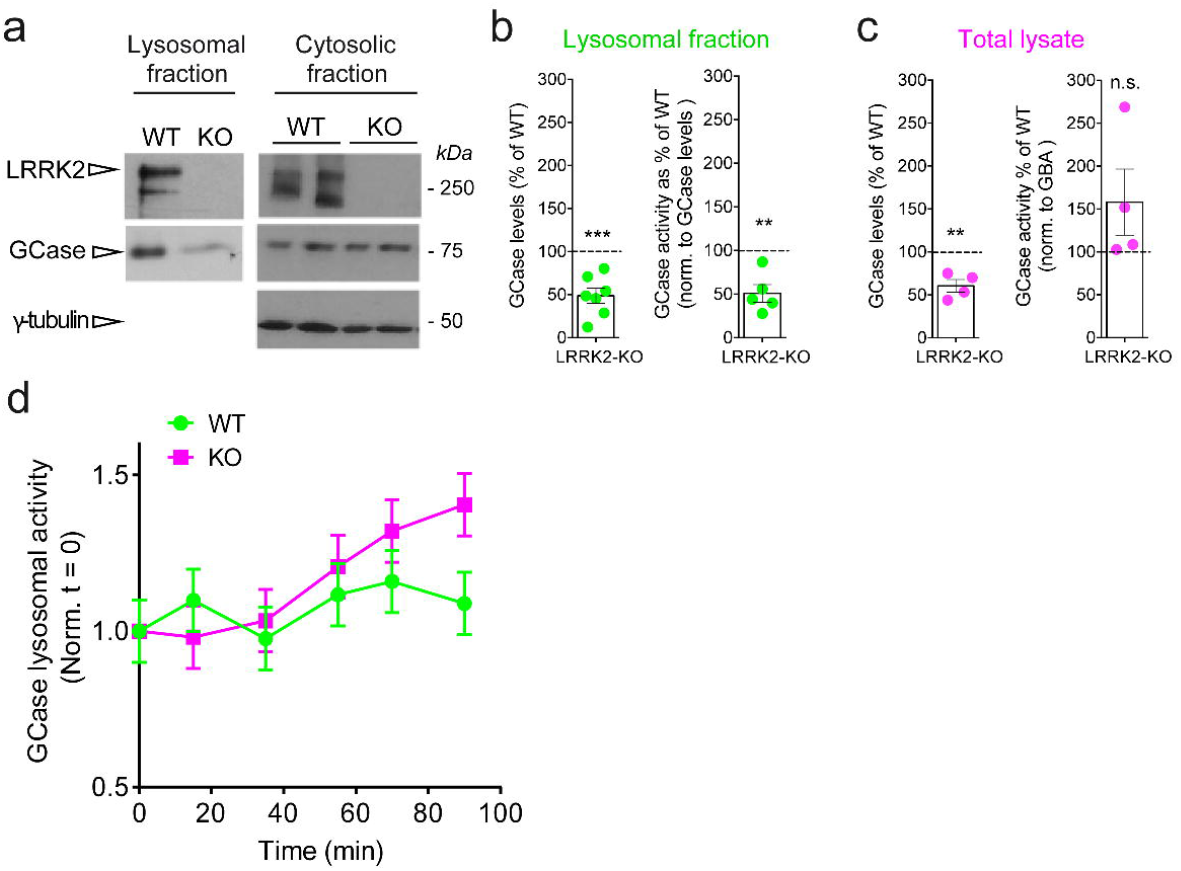
GCase activity and level is reduced in lysosomal extracts from LRRK2 KO RAW264.7 cells. a. RAW264.7 cells, WT and LRRK2-deficient (KO) were processed for western immunoblotting of the cytoplasmic fraction and crude lysosomal fraction, and the membranes probed for LRRK2, GCase, and γ-tubulin. GCase levels are unchanged in the cytoplasmic fraction of LRRK2 KO cells, compared to WT cells; however, are reduced in crude lysosomal fractions from KO RAW264.7 cells. b. In the lysosomal fraction of RAW264.7 cells, levels of GCase are reduced by approximately 50% in LRRK2-KO cells, compared to WT (set to 100%); and the activity of GCase, normalized to GCase expression levels, is also significantly reduced. (Two-tailed t-tests were performed; ** p<0.01, *** p<0.001). c. In total cell extracts of RAW264.7 cells, GCase expression levels are likewise reduced in KO cells compared to WT (set to 100%) cells; whereas normalized GCase activity showed a non-significant increase compared to WT cells. (Two-tailed t-tests were performed; ** p<0.01). d. Representative graph (n=3 technical replicates) of lysosomal GCase activity measured over time in live cells using the PDFGlu probe and normalized by the total protein level at the final time point. These findings suggest that upon MLi-2 treatment, lysosomal GCase activity in WT RAW264.7 cells is increased to the level observed for LRRK2 KO macrophages. However, upon MLi-2 treatment LRRK2 KO RAW264.7 cells show an increase in lysosomal GCase activity, which would suggest a spurious effect of MLi-2 on GCase that is not mediated by LRRK2. This trend was observed in 3 out of 4 independent experiments with identical setup.

As the 4-MU substrate-based assay allows us to assess the intrinsic activity of GCase outside its native cellular context, we were prompted to investigate GCase activity under native conditions using the live cell-based fluorescent probe PFB-FDGlu. Following published methods^16^, we first pre-incubated the cells with the GCase probe for 60 minutes to allow its internalization and then measured the fluorescence increase over time. As shown in figure 6c, there was a modest increase over time in GCase activity in KO cells compared with WT cells upon normalization to total protein levels (Fig. 6d).

Overall, these results indicate that: 1) GCase activity measured in cells differs from GCase activity probed *in vitro*; 2) the low in-cell GCase activity measured in WT cells may reflect a faster substrate consumption as compared with KO cells occurring within the first 60 minutes of dye uptake^27^; 3) alteration in the lysosomal pH may affect the assay outcome, since PFB-FDGlu can be a pH-sensitive dye and LRRK2 mutants are known to alter pH in the endolysosomal compartment in certain cell types^28,29^.

## Discussion

In this work we showed that an overall positive correlation exists between LRRK2 kinase activity and GCase hydrolytic activity *ex-vivo* using a wide range of PD-relevant cellular and tissue models, as well as human clinical samples and patient-derived iPSC dopaminergic neurons. Our assessments reflect the intrinsic activity of GCase outside the cellular context upon normalization by GCase protein levels (when possible). Interestingly, GCase protein levels also appear to be negatively affected by LRRK2 kinase activity and expression; yet, when normalized by GCase protein levels, the intrinsic activity of the enzyme is increased in the context of *LRRK2* GSKI brains. Interestingly, GCase protein level is reduced also in the cytosolic and lysosomal fraction of extracts of LRRK2 KO macrophages. These results are in agreement with our previous study where we detected reduced GCase protein levels in the whole brain of 12 month old LRRK2 KO mice and increased GCase activity when normalized by GCase levels^15^. Recently, the group of Morari and colleagues failed to detect a difference in striatal GCase expression levels in three or 12 month old WT and KO mice^30^; however, striatal GCase activity was reported to be increased in KO and kinase dead KI mice. In contrast, G2019S-KI mice did not exhibit differences in striatal GCase activity. The difference in findings between this report, our earlier study in vivo, and the current study are likely due to methodological differences, such as the normalization of GCase activity and the use of a specific GCase inhibitor (e.g. CBE) to reveal changes in specific activity.

In contrast to what was observed in total lysates, crude lysosomal preparations from LRRK2 KO macrophages display decreased intrinsic GCase activity. One possible interpretation is that the amount of GCase that is correctly trafficked to the lysosomes in RAW LRRK2 KO is less, and also less active, while the overall GCase produced in cells may still be reduced but its activity would not be affected before it is trafficked to the lysosomes. Another possibility is that the increased protein degradation generally observed in LRRK2 KO^31^ affects the amount of GCase in the lysosomes. We were not able to reliably measure specific GCase activity in the resulting cytoplasmic fraction, remaining after the gradient enrichment of lysosomes, but the measurements obtained from total lysates indicate this as a possible scenario.

To further support that the alterations in GCase are correlated to LRRK2 kinase activity, we investigated GCase level and activity in HEK293T overexpressing LRRK2 G2019S. Two approaches were used to manipulate LRRK2 kinase activity and monitor the relative GCase function in these models. A genetic approach expressing a kinase-dead/G2019S LRRK2 double mutant (G2019S/K1906R) in HEK293T cells was chosen to confirm the results obtained via pharmacological kinase inhibition. Consistently, these experiments support the positive correlation between GCase hydrolytic activity and LRRK2 kinase activity *in vitro*.

In addition to these cellular models, we assessed GCase activity in additional patient-derived models more relevant for LRRK2-PD pathology: PBMCs, plasma, fibroblasts and iPSC-derived neurons. Overall, a positive correlation between LRRK2 kinase and GCase hydrolytic activities is observed in all these systems, although to different extents. In PBMCs, *LRRK2* G2019S PD patients display an increased GCase activity compared with healthy controls^14^, while the activity of GCase in PBMCs from non-manifesting *LRRK2* G2019S carriers is similar to that of controls. This may suggest that a synergistic effect of the mutation and of the disease status occurs, at least in this immune cell population. Another possible explanation is that GCase activity alterations in subjects carrying *LRRK2* G2019S mutations are measurable in this cell type only when the disease is already clinically manifested, suggesting that GCase activity in these patients may be used as a marker for disease progression or manifestation. Of note, a similar trend is observed in plasma, where manifesting *LRRK2* G2019S carriers exhibit higher (although not significant) GCase activity as compared with iPD or healthy controls. It has been previously reported that lysosomal enzyme activity can be measured in the plasma of patients^32^, suggesting that these proteins are likely to be secreted in certain conditions. In particular, this may occur via lysosomal exocytosis, which is a Ca^2+^-regulated process important for plasma membrane repair and secretion^33^. This may be relevant and deserves further investigation to understand if LRRK2 is involved in the regulation of the pathway and to evaluate if GCase measurement in the plasma (together or alternatively to PBMCs) could be used as a biomarker at least for certain PD forms and may allow patient stratification. Given the participation of LRRK2 in such pathways, it is reasonable to envision a role for LRRK2 in regulating GCase release or secretion.

GCase activity is significantly reduced in *LRRK2* G2019S gene-corrected iPSC-derived neurons, whereas it is increased in control iPSC-derived neurons in which the G2019S mutation has been introduced. This suggests that, at least in this cell type, increased LRRK2 kinase activity is itself sufficient to impact on GCase and may be a peculiar feature of dopaminergic neurons, associated with their prominent vulnerability to cell death in PD. In contrast to our current and previous findings showing no difference in activity in mixed PBMCs from two independent iPD cohorts^20^, the group of Dzamko reports reduced GCase activity in isolated CD14+ monocytes from idiopathic PD patients^11^. A major difference from the two studies is that Atashrazm and co-authors used an in-cell (PFB-FDGlu) method for assessing GCase activity, but also raises the possibility that cell-type specific alterations in GCase activity may exist in the different immune cell sub-types, since PBMCs are a more heterogeneous cell population than CD14+ monocytes. Further studies assessing GCase activity (using both methodologies) in purified monocytes from LRRK2 mutation carriers, are warranted. Our finding of decreased GCase activity in PD-patients carrying *GBA1* mutations is consistent with previous reports in other models^9,34,35^ and suggests that impaired GCase activity may be a general contributing factor to PD, yet clearly interacting with certain genetic backgrounds.

It has been reported that deficiency in GCase activity leads to the accumulation of α-synuclein^36^. The increased GCase activity in *LRRK2* G2019S patients may explain, at least in part, the lack of Lewy Bodies in a subpopulation of LRRK2-patients^37^, a hypothesis that requires further investigations. In the case of *LRRK2* G2019S PBMCs and iPSC-derived neurons, normalization by GCase expression levels could not be performed. However, if the trend in GCase protein levels is consistent with the other models, GCase intrinsic activity may further increase also in these systems.

In human fibroblasts, as for all other human-derived cell types, the interindividual variability was even more pronounced, and despite a trend for increase in GCase activity in mutant LRRK2-patient derived fibroblasts, this did not reach statistical significance. What is striking is instead the presence of MLBs in the *LRRK2* G2019S cells that are almost completely absent in control fibroblasts. The number of MLBs is negatively correlated with the GCase activity when normalized by GCase level: the lower the activity, the more MLBs are present. This resembles the behavior of fibroblasts from PD patients carrying the N370S *GBA1* mutation, which have reduced GCase activity and accumulate MLBs^21^. This may indicate that LRRK2 mutant-expressing cells that fail to mount an adequate response via upregulation of GCase activity transition to a state of dysfunction of the ALP.

Technical differences in the methods used to measure GCase activity and to normalize it may account for the different results reported by Krainc and Dzamko groups^11,16^. In those studies, lysosomal GCase activity was measured in iPSC-derived *LRRK2* G2019S neurons using the live-cell probe PFB-FDGlu rather than GCase activity *in vitro*, which is more akin to assess the intrinsic activity of the isolated enzyme, possibly reflecting certain post-translational modifications of the protein itself. In contrast, a cellular-based GCase probe would not just reflect the net effect of the intrinsic GCase activity, but also the activities of other lysosomal enzymes and co-factors, the trafficking of GCase to the lysosome, and lysosomal pH. It should also be noted that the cellular GCase probe utilized in these studies may also be sensitive to lysosomal pH, which is known to be altered in the context of *LRRK2* G2019S expression^28,29^. Additionally, the cellular uptake of this probe, through pinocytosis, may also be affected by LRRK2 activity^27^, further complicating the interpretation. Here, we also attempted to measure in-cell GCase activity using the PFB-FDGlu live-cell probe and found that LRRK2 KO macrophages display a trend for an increase in GCase activity (Fig. 6), a result in agreement with the previous studies^11,16^. However, we feel that caution should be taken in interpreting these data as we do not have information on GCase activity within the first 60 minutes of probe uptake. This timeframe is required to allow substrate internalization but, clearly, the substrate will be converted into its reaction product starting from t=0. Thus, the ratio between internalized over converted probes may be a more representative, though technically complicated, measurement.

The mechanisms by which LRRK2 kinase activity mediates GCase function seems to be related to both GCase level and its intrinsic activity. Given its prominent role in membrane trafficking, it is tempting to speculate that one potential mechanism for the effects of LRRK2 activity on GCase function is due to altered trafficking to the lysosome. Accordingly, in our enriched lysosomal preps, we detect a significant increase in GCase activity *in vitro* in the presence of *LRRK2* G2019S. To determine whether the impact of LRRK2 on lysosomal GCase content and activity is specific, it will be critical to evaluate whether GCase transporters or co-factors, such as LIMP2 and the M6P receptor, or other lysosomal proteins are affected in LRRK2 models.

Finally, carriers of both *GBA1* and *LRRK2* mutations are frequently reported, particularly within specific ethnic groups such as the Ashkenazi-Jewish community. While clinical reports in the literature of such double carriers are few, in one study it was found that the G2019S mutation in mutant *GBA1* carriers appeared to reduce the likelihood of patients developing non motor symptoms^38^. In another recent study, *LRRK2* G2019S/*GBA1* PD patients showed a more benign course of the disease with respect to both motor and non-motor symptoms when compared to *GBA1*-PD patients^39^. Our findings in this study point to the possibility that this is due to the balance between the G2019S mutation in LRRK2, which increases GCase activity at least in certain conditions, and *GBA1* mutations that reduced GCase enzymatic function. However, this will need to be experimentally proven.

Clearly, more studies are warranted to uncover the link between these two key players in PD pathogenesis and progression and the precise molecular mechanisms by which this interplay occurs in the different idiopathic and genetic forms of the disease; and importantly, to have a better understanding of the functional assessment of GCase activity (*in vitro* vs whole cell) and its interpretation.

## Materials and Methods

### Patient demographics

The samples assessed in this study were collected from patients and control subjects at the Movement Disorder division in the Department of Neurology at Columbia University Irving Medical Center. The demographic and clinical data of the participants have been described in more detail elsewhere^19^. Participants were screened for the *LRRK2* G2019S mutation, as well as several *GBA1* mutations and variants (see Table 1). All samples were collected under informed consent, and the study protocol was approved by the IRBs of both Columbia University (CUIMC) as well as the Biomedical Research Foundation of the Academy of Athens (BRFAA).

### PBMC isolation

PBMCs were isolated using standard protocols. Briefly, sodium citrate Vacutainer tubes (BD) were used to collect whole blood from study participants. Blood was diluted 1X in sterile PBS and transferred to Ficoll-containing Leucosep tubes (Griener) and centrifuged at 1000xg for 10 minutes at room temperature. The upper plasma phase was then extracted and centrifuged at 300xg for 15 minutes at room temperature. The banded cells were collected and washed 1X in PBS and centrifuged again at 300xg for 10 minutes before counting and aliquoting, in RPMI with 40% FBS and 10% DMSO, at 3 × 106 viable cells per cryovial. Frozen cells were stored at -80ºC. In total, we report results from 70 participants, including 20 idiopathic PD, 18 healthy controls, 10 GBA1-PD, 4 non-manifesting GBA1, 8 LRRK2-PD, 9 non-manifesting LRRK2, and one subject with PD and both GBA1 and LRRK2.

### Plasma isolation

Whole blood was collected in EDTA vacutainer tubes, which were centrifuged at 1,500xg for 15 minutes at 4º C, from which 1mL plasma aliquots were extracted within 60 minutes of collection. Samples were stored in a -80°C freezer until processing. In total, we report results from 40 subjects, including 10 idiopathic PD, 10 healthy controls, 10 GBA1-PD, and 10 LRRK2-PD subjects.

### Cell lines, plasmids, and transfection

Human HEK293T cells and murine RAW264.7 cells control and Lrrk2 knockouts (ATCC), were grown in DMEM (Sigma; high glucose) supplemented with 10% FBS and penicillin/streptomycin. Primary fibroblasts, obtained from the Coriel Institute and from Istituto G. Gaslini^40^, were grown in DMEM plus 10% FBS and antibiotics. Plasmids encoding Flag-tagged wild type or mutant LRRK2 were previously described^41,42^. For transient transfection, HEK293T cells were incubated with DNA:CaPO_4_ precipitates or PEI as transfection reagents, and growth media was changed 24h later.

### Crude lysosomal isolation

HEK293T or RAW264.7 cells were washed once with PBS, then collected in PBS and centrifuged at 3500 rpm for 5 min at 4°C. The cell pellets were lysed in 0.25M Sucrose/1xTBS buffer (pH 7.4) supplemented with 1x phosphatase inhibitors (Roche) in a manual glass homogenizer. The lysates were centrifuged at 6,800 x g for 15min at 4°C. The supernatant was kept and the pellet was then washed with the same sucrose lysis buffer and centrifuged again as before. The combined supernatants from the previous steps were centrifuged at 21,000 x g for 30min at 4°C to yield a pellet containing lysosomes and supernatant containing cytoplasmic proteins. The lysosomal-enriched pellet was washed with the sucrose buffer and then resuspended in 50μl activity assay buffer (50mM Citric acid, 176mM K2HPO4, 10mM Sodium taurocholate, 0.01% Tween-20, pH 5.9). The lysosomal lysates were kept at -80°C or were processed for analysis immediately.

### Electron microscopy of human fibroblasts

Samples were fixed with 2.5% glutaraldehyde in 0.1 M sodium cacodylate buffer pH 7.4 ON at 4°C. The samples were postfixed with 1% osmium tetroxide plus potassium ferrocyanide 1% in 0.1M sodium cacodylate buffer for 1 hour at 4°. After three water washes, samples were dehydrated in a graded ethanol series and embedded in an epoxy resin (Sigma-Aldrich). Ultrathin sections (60-70nm) were obtained with an Ultrotome V (LKB) ultramicrotome, counterstained with uranyl acetate and lead citrate and viewed with a Tecnai G2 (FEI) transmission electron microscope operating at 100 kV. Images were captured with a Veleta (Olympus Soft Imaging System) digital camera.

### Human differentiated dopaminergic neurons

Fibroblasts were isolated from skin biopsies of 4 control subjects with wild type *LRRK2*, 2 unrelated Parkinson’s disease patients carrying a heterozygous G2019S mutation of the *LRRK2* gene, 1 Parkinson’s disease patient carrying a heterozygous L444P mutation of *GBA1* gene, and 2 Parkinson’s disease patients carrying *SNCA* gene mutations (gene duplication and the heterozygous A53T mutation). Fibroblasts were reprogrammed into iPSCs through viral transfection of Klf4–Oct3/4–Sox2, cMyc, and Klf4 (CytoTune™-iPS 2.0 Sendai Reprogramming Kit, ThermoFisher Scientific). One of the *LRRK2*-mutated iPSC line underwent genome editing to obtain an isogenic control with corrected *LRRK2* (LRRK2 #2 GC). In the same way, another control iPSC line (CTR5) was genome edited to generate a mutant G2019S *LRRK2* line (CTR5 G2019S). iPSCs were cultured in E8 medium (ThermoFisher Scientific) and passaged with EDTA 5 mM every 1-3 days. Generated iPSCs underwent karyotype analysis, which was negative for chromosomal rearrangements. Proper reprogramming of iPSCs was assessed by expression of stem cell markers (SOX2, OCT4, TRA-1-60, SSEA4).

All iPSC lines were differentiated into dopaminergic neurons according to the protocol described by Kriks et al. (2011)^43^ with modifications as described in Monzio Compagnoni et al. (2018)^44^. Immunofluorescent analysis for markers of dopaminergic identity (TUJ1, TH, and DAT) was performed to assess dopaminergic differentiation. Antibodies were used with following dilutions: TUJ1 (Abcam ab18207, 1:250), TH (Thermo Scientific PA5-17800 1:100), DAT (Millipore Ab2231, 1:600).

### LRRK2 transgenic animals

Homozygous LRRK2 G2019S (GSKI) and WT mice were housed at the National Institute on Aging, NIH, according to a protocol approved by the Institutional Animal Care and Use Committee of the National Institute on Aging, NIH (463-LNG-2019). Dissections of cortex, midbrain and striatal regions were performed in 6 months-old mice of all genotypes. 4-5 mice/genotype were used in all experiments.

### Brain and fibroblast lysates preparation

Brain lysates were obtained by mechanically lysing the different regions on ice in 25 mM pH 7.5 Tris-HCl, 150 mM NaCl, 1% (v/v) NP40, 1% (w/v) sodium deoxycholate, 0,1% (w/v) SDS, 2 mM EGTA, 20 mM sodium fluoride, 50mM beta glycerophosphate, 50 mM sodium pyrophosphate, 20 mM sodium orthovanadate. Fibroblasts were lysed in RIPA buffer, supplemented with protease inhibitors. Protein lysates were clarified by centrifugation and total protein concentration was determined by BCA assay (PierceTM, Thermo Fisher). Fifty μg of protein samples for brain lysates and 20 μg of protein samples were resolved on polyacrylamide gels (see Western Immunolotting for details).

### Measurement of GCase activity

We assessed activity of β-glucocerebrosidase (GCase) using two different approaches. With the fluorescent GCase substrates blue fluorogenic substrate 4-methylumbelliferyl-β-D-glucopyranoside (4-MU) (Sigma-Aldrich; as described in ^11,12^), PBMCs were washed 2x with PBS and lysed in 60μl of activity assay buffer (50mM Citric acid, 176mM K_2_HPO_4_, 10mM Sodium taurocholate, 0.01% Tween-20, pH 5.9) and we used for this assay a standard volume of 5μl from total lysates. These total or lysosomal lysates were diluted in 5μl activity assay buffer (see above) or 5μl of 20mM Conduritol B Epoxide (CBE) (Santa Cruz, sc-201356A) (specific GCase inhibitor) and incubated for 15 min at RT. Subsequently, 25μl of 5mM 4-MU (Sigma, M3633) were added and immediately incubated for 25 min at 37°C. The reaction was stopped by adding 465μl stop buffer (1M NaOH, 1M Glycine, pH 10). Fluorescence was measured (excitation 360 nm, emission 450 nm) in glass cuvettes in luminescence spectrometer (Perkin Elmer LS 55), given as relative fluorescence units (RFU). Each reaction was performed in duplicate. For each sample, the mean values in the presence or absence of the CBE inhibitor were calculated. All measurements were corrected by subtracting the average of nonspecific activity (incubation with CBE inhibitor) from specific average activity (incubation with activity assay buffer). The resulting measurements were then normalized to band intensities of GCase protein for lysosomal fractions of cell lines. For PBMCs the GCase activity measurements were first normalized to total protein concentration (Bradford method); or, as an alternative, to the band intensities of the GBA/GAPDH ratio following Western immunoblotting using ImageJ.

GCase activity was also measured in lysates from brain regions (midbrain, cortex and striatum) of different transgenic mice, lysed as previously described, and in lysates from human fibroblasts. Brain or fibroblast lysates were diluted in McIlvaine buffer pH 4.5 (standard volume of 2-5 µl from total lysates was used), and a final concentration of 2mM of the fluorogenic substrate 4MU (Sigma, M3633) were added to reach a final volume of 60µl and immediately incubated for 90 min at 37°C. The reaction was stopped by adding 200μl of stop buffer (1M NaOH, 1M Glycine, pH 10). Fluorescence was measured (ex 360 nm, em 450 nm) in a plate reader (Victor X3, Perkin Elmer). Three technical replicates were performed for each experiment and all measurements were corrected by background subtraction. The resulting measurements were either normalized to band intensities of total GCase protein or by the total proteins in solution (and this is clearly stated in the results). A similar protocol was used to measure GCase activity in plasma from PD patients and healthy controls. In this experiment, the enzymatic activity assay was performed on 20μl of plasma and normalized by total protein concentration as evaluated via BCA assay.

For GCase activity detection in dopaminergic neurons, 20 µg of proteins were incubated for 30 minutes at room temperature with 25 µl of McIlvaine buffer 4X (0.4 M citric acid, 0.8 M Na2HPO4) pH 5.2, AMP-DNM (N-(5-adamantane-1-yl-methoxy-pentyl)-Deoxynojirimycin) at a final concentration of 5 nM, and H2O to a final volume of 100 µl. At the end of incubation, 25 µl of 4-MU was added at a final concentration of 6 mM and incubated for one hour at 37°C. At the end of incubation, 10 µl of the reaction mixture were transferred to black 96-well plates and 190 µl of 0.25 M glycine pH 10.7 were added. Plates were read by Victor microplate reader (Perkin Elmer). GCase activity was expressed as picomoles of converted substrate/mg of proteins/minute.

The second approach to assess GCase activity is based on the measurement of cellular fluorescence of the Fluorescein di-β-D-glucopyranoside (PFB-FDGlu) (Marker gene technologies) probe in a fluorescence microplate reader (Victor X3, Perkin Elmer). Briefly, the cells were incubated with 50μg/ml of PFB-FDGlu for 60 min at 37ºC. Parallel cells were pre-treated with 100 μM CBE (Santa Cruz,) for 1hr at 37ºC before adding the PFB-FDGlu substrate. Specific GCase activity was expressed as the difference in fluorescence signal with and without treatment with CBE.

### Western Immunoblotting

Equal amounts of protein were loaded on 10% SDS-PAGE gels. For brain lysates, 18-well Criterion SDS-gels were used (Cat# 1704273, Biorad). Primary antibodies were diluted in 5% milk/TBST or 5% BSA/TBST as recommended, and incubated with first antibodies overnight at 4°C. The following primary antibodies were used: rabbit anti-LRRK2 (abcam, UDD3 ab170993, 1:2000), rabbit anti-pS1292-LRRK2 (abcam, ab206035, 1:2000), rabbit anti-pS935-LRRK2 [UDD2] (abcam, ab172382, 1:1000), rabbit anti-LRRK2 (abcam, c41-2 ab133474, 1:1000), rabbit anti-Glucocerebrosidase (Sigma, G4171, 1:1000), rabbit anti-Rab29 (abcam, ab199644, 1:1000), rabbit anti-pT71-Rab29 (abcam, ab241062), mouse anti-γ tubulin (Sigma, T5326-25UL), mouse anti-GAPDH (Millipore, #MAB374), rabbit anti-LIMP2 (Thermo Fisher Scientific, AB_2182836, 1:1000), rabbit anti-SOD1 (Santa Cruz, sc11407, 1:1000), rabbit anti-human LAMP2A (abcam, ab23322, 1:1000), mouse anti-LAMP1 (sc-20011). After washing 3 times for 10min with TBST, the membranes were incubated with the appropriate secondary antibody conjugated with horseradish-peroxidase (HRP). Bands were visualized with chemiluminescence using Clarity ECL substrate (Biorad) or Immobilon Forte Western HRP Substrate (WBLUF0100, Merck-Millipore) as per manufacturer’s instructions.

### Statistical analyses

All quantitative data are expressed as mean ± SEM (standard error of the mean) from at least 4 different mice/genotype or at least 3 independent cell cultures. Significance of differences between two groups was verified by Student t-test while comparisons between 3 or more groups were performed by one-way ANOVA with Dunnett’s Multiple comparison test/Bonferroni’s post-hoc test/Tukey’s post-hoc test.

## Acknowledgements

We thank Michael J Fox Foundation for the support to this research. NP postdoctoral fellowship was funded by Fondazione Veronesi (2018-2019). We also thank the “Cell line and DNA biobank from patients affected by genetic diseases” (Istituto G. Gaslini), and “Parkinson Institute Biobank” (Milan, http://www.parkinsonbiobank.com/) members of the Telethon Network of Genetic Biobanks funded by Telethon Italy, (http://www.biobanknetwork.org, project No. GTB12001). We acknowledge the Imaging Facility at the Department of Biology, University of Padova (Italy).

## Author contributions

M. K. performed the experiments on PBMCs and part of the experiments on RAW and HEK cells. E. F. and P. B. prepared and characterized the iPSC-derived neurons and M. A. performed the biochemical analysis on iPSC-derived neurons.

E. Z. performed TEM experiments. S. C. and N. P. performed the experiments on mice brains, and part of the experiments on RAW and HEK cells. A. K. handled the mice colony and collected brain samples. M. D. and A. D. F. supervised the iPSC work. L. C contributed to data analysis and discussion. M. S. and R. N. A. selected the patient cohort for the PMBCs and plasma experiments and collected the samples. M. C. and L. S. provided data interpretation. E. G., H. R. and N. P. conceived the experimental plan, supervised the work and wrote the manuscript. All the authors contributed to the manuscript preparation.

## Competing interests

The authors declare no competing interests.

## Notes

### Competing Interest Statement

The authors have declared no competing interest.

## References

1 Paisán-Ruíz C, Jain S, Evans EW, Gilks WP, Simón J, Van Der Brug M et al. Cloning of the gene containing mutations that cause PARK8-linked Parkinson’s disease. Neuron 2004; 44: 595–600.

2 Nalls MA, Blauwendraat C, Vallerga CL, Heilbron K, Bandres-Ciga S, Chang D et al. Identification of novel risk loci, causal insights, and heritable risk for Parkinson’s disease: a meta-analysis of genome-wide association studies. Lancet Neurol 2019; 18: 1091–1102.

3 Zimprich A, Biskup S, Leitner P, Lichtner P, Farrer M, Lincoln S et al. Mutations in LRRK2 cause autosomal-dominant parkinsonism with pleomorphic pathology. Neuron 2004; 44: 601–607.

4 Sidransky E, Nalls MA, Aasly JO, Aharon-Peretz J, Annesi G, Barbosa ER et al. Multicenter Analysis of Glucocerebrosidase Mutations in Parkinson’s Disease. N Engl J Med 2009; 361: 1651–1661.

5 Sidransky E, Lopez G. The link between the GBA gene and parkinsonism. Lancet Neurol 2012; 11: 986–998.

6 Bonet-Ponce L, Cookson MR. LRRK2 recruitment, activity, and function in organelles. FEBS J 2021;:1–20.

7 Steger M, Tonelli F, Ito G, Davies P, Trost M, Vetter M et al. Phosphoproteomics reveals that Parkinson’s disease kinase LRRK2 regulates a subset of Rab GTPases. Elife 2016; 5: 1–28.

8 Di Maio R, Hoffman EK, Rocha EM, Keeney MT, Sanders LH, De Miranda BR et al. LRRK2 activation in idiopathic Parkinson’s disease. Sci Transl Med 2018; 10. doi:10.1126/scitranslmed.aar5429.

9 Malini E, Grossi S, Deganuto M, Rosano C, Parini R, Dominisini S et al. Functional analysis of 11 novel GBA alleles. Eur J Hum Genet 2014; 22: 511–516.

10 Mazzulli JR, Xu YH, Sun Y, Knight AL, McLean PJ, Caldwell G a. et al. Gaucher disease glucocerebrosidase and α-synuclein form a bidirectional pathogenic loop in synucleinopathies. Cell 2011; 146: 37–52.

11 Atashrazm F, Hammond D, Perera G, Dobson-Stone C, Mueller N, Pickford R et al. Reduced glucocerebrosidase activity in monocytes from patients with Parkinson’s disease. Sci Rep 2018; 8: 1–12.

12 Hughes LP, Pereira MMM, Hammond DA, Kwok JB, Halliday GM, Lewis SJG et al. Glucocerebrosidase Activity is Reduced in Cryopreserved Parkinson’s Disease Patient Monocytes and Inversely Correlates with Motor Severity. J Parkinsons Dis 2021; pre-press: 1–9.

13 Gegg ME, Burke D, Heales SJR, Cooper JM, Hardy J, Wood NW et al. Glucocerebrosidase deficiency in substantia nigra of parkinson disease brains. Ann Neurol 2012; 72: 455–463.

14 Alcalay RN, Levy OA, Waters CC, Fahn S, Ford B, Kuo SH et al. Glucocerebrosidase activity in Parkinson’s disease with and without GBA mutations. Brain 2015; 138: 2648–2658.

15 Ferrazza R, Cogo S, Melrose H, Bubacco L, Greggio E, Guella G et al. LRRK2 deficiency impacts ceramide metabolism in brain. Biochem Biophys Res Commun 2016; 478. doi:10.1016/j.bbrc.2016.08.082.

16 Ysselstein D, Nguyen M, Young TJ, Severino A, Schwake M, Merchant K et al. LRRK2 kinase activity regulates lysosomal glucocerebrosidase in neurons derived from Parkinson’s disease patients. Nat Commun 2019; 10: 1–9.

17 Sosero YL, Yu E, Krohn L, Rudakou U, Mufti K, Ruskey JA et al. LRRK2 p.M1646T is associated with glucocerebrosidase activity and with Parkinson’s disease. Neurobiol Aging 2021; 103: 142.e1-142.e5.

18 Sanyal A, DeAndrade MP, Novis HS, Lin S, Chang J, Lengacher N et al. Lysosome and Inflammatory Defects in GBA1-Mutant Astrocytes Are Normalized by LRRK2 Inhibition. Mov Disord 2020; 35: 760–773.

19 Melachroinou K, Kang MS, Liong C, Narayan S, Levers N, Joshi N et al. Elevated In Vitro Kinase Activity in Peripheral Blood Mononuclear Cells of Leucine-Rich Repeat Kinase 2 G2019S Carriers: A Novel Enzyme-Linked Immunosorbent Assay–Based Method. Mov Disord 2020; 35: 2095–2100.

20 Papagiannakis N, Xilouri M, Koros C, Stamelou M, Antonelou R, Maniati M et al. Lysosomal alterations in peripheral blood mononuclear cells of Parkinson’s disease patients. Mov Disord 2015; 30: 1830–1834.

21 García-Sanz P, Orgaz L, Bueno-Gil G, Espadas I, Rodríguez-Traver E, Kulisevsky J et al. N370S-GBA1 mutation causes lysosomal cholesterol accumulation in Parkinson’s disease. Mov Disord 2017; 32: 1409–1422.

22 García-Sanz P, Orgaz L, Fuentes JM, Vicario C, Moratalla R. Cholesterol and multilamellar bodies: Lysosomal dysfunction in GBA-Parkinson disease. Autophagy 2018; 8627: 1–2.

23 Sakurai M, Kuwahara T. Two Methods to Analyze LRRK2 Functions Under Lysosomal Stress: The Measurements of Cathepsin Release and Lysosomal Enlargement. In: Experimental Models of Parkinson’s Disease. 2021, pp 63–72.

24 Nazish I, Arber C, Piers TM, Warner TT, Hardy JA, Lewis PA et al. Abrogation of LRRK2 dependent Rab10 phosphorylation with TLR4 activation and alterations in evoked cytokine release in immune cells. Neurochem Int 2021; 147: 105070.

25 Eguchi T, Kuwahara T, Sakurai M, Komori T, Fujimoto T, Ito G. LRRK2 and its substrate Rab GTPases are sequentially targeted onto stressed lysosomes and maintain their homeostasis. Proc Natl Acad Sci 2018; 115: E9115–E9124.

26 Herbst S, Campbell P, Harvey J, Bernard EM, Papayannopoulos V, Wood NW et al. LRRK 2 activation controls the repair of damaged endomembranes in macrophages. EMBO J 2020; 44: 1–14.

27 Liu Z, Xu E, Zhao HT, Cole T, West AB. LRRK2 and Rab10 coordinate macropinocytosis to mediate immunological responses in phagocytes. EMBO J 2020; 39: 1–21.

28 Henry AG, Aghamohammadzadeh S, Samaroo H, Chen Y, Mou K, Needle E et al. Pathogenic LRRK2 mutations, through increased kinase activity, produce enlarged lysosomes with reduced degradative capacity and increase ATP13A2 expression. Hum Mol Genet 2015; 24: 6013–6028.

29 Madureira M, Connor-Robson N, Wade-Martins R. “LRRK2: Autophagy and Lysosomal Activity”. Front Neurosci 2020; 14. doi:10.3389/fnins.2020.00498.

30 Albanese F, Mercatelli D, Finetti L, Lamonaca G, Pizzi S, Shimshek DR et al. Constitutive silencing of LRRK2 kinase activity leads to early glucocerebrosidase deregulation and late impairment of autophagy in vivo. Neurobiol Dis 2021; 159: 105487.

31 Pellegrini L, Hauser DN, Li Y, Mamais A, Beilina A, Kumaran R et al. Proteomic analysis reveals co-ordinated alterations in protein synthesis and degradation pathways in LRRK2 knockout mice. Hum Mol Genet 2018; 27: 3257–3271.

32 Ungewickell AJ, Majerus PW. Increased levels of plasma lysosomal enzymes in patients with Lowe syndrome. Proc Natl Acad Sci U S A 1999; 96: 13342–13344.

33 Tancini B, Buratta S, Delo F, Sagini K, Chiaradia E, Pellegrino RM et al. Lysosomal exocytosis: The extracellular role of an intracellular organelle. Membranes (Basel) 2020; 10: 1–21.

34 Ron I, Horowitz M. ER retention and degradation as the molecular basis underlying Gaucher disease heterogeneity. Hum Mol Genet 2005; 14: 2387–2398.

35 Fernandes HJR, Hartfield EM, Christian HC, Emmanoulidou E, Zheng Y, Booth H et al. ER Stress and Autophagic Perturbations Lead to Elevated Extracellular α-Synuclein in GBA-N370S Parkinson’s iPSC-Derived Dopamine Neurons. Stem Cell Reports 2016; 6: 342–356.

36 Du TT, Wang L, Duan CL, Lu LL, Zhang JL, Gao G et al. GBA deficiency promotes SNCA/alpha-synuclein accumulation through autophagic inhibition by inactivated PPP2A. Autophagy 2015; 11: 1803–1820.

37 O’Hara DM, Pawar G, Kalia SK, Kalia L V. LRRK2 and α-Synuclein: Distinct or Synergistic Players in Parkinson’s Disease? Front Neurosci 2020; 14. doi:10.3389/fnins.2020.00577.

38 Yahalom G, Rigbi A, Israeli-Korn S, Krohn L, Rudakou U, Ruskey J et al. Age at Onset of Parkinson’s Disease Among Ashkenazi Jewish Patients: Contribution of Environmental Factors, LRRK2 p.G2019S and GBA p.N370S Mutations. J Park s Dis 2020; 3: 1123–1132.

39 Omer N, Giladi N, Gurevich T, Bar-Shira A, Gana-Weisz M, Goldstein O et al. A Possible Modifying Effect of the G2019S Mutation in the LRRK2 Gene on GBA Parkinson’s Disease. Mov Disord 2020; 35: 1249–1253.

40 Filocamo M, Mazzotti R, Corsolini F, Stroppiano M, Stroppiana G, Grossi S et al. Cell Line and DNA Biobank From Patients Affected by Genetic Diseases. Open J Bioresour 2014; 1: e2.

41 Civiero L, Cogo S, Kiekens A, Morganti C, Tessari I, Lobbestael E et al. PAK6 Phosphorylates 14-3-3 γ to Regulate Steady State Phosphorylation of LRRK2. Front Mol Neurosci 2017; 10. doi:10.3389/fnmol.2017.00417.

42 Ho CCY, Rideout HJ, Ribe E, Troy CM, Dauer WT. The Parkinson disease protein leucine-rich repeat kinase 2 transduces death signals via Fas-associated protein with death domain and caspase-8 in a cellular model of neurodegeneration. J Neurosci 2009; 29: 1011–1016.

43 Kriks S, Shim J-W, Piao J, Ganat YM, Wakeman DR, Xie Z et al. Dopamine neurons derived from human ES cells efficiently engraft in animal models of Parkinson’s disease. Nature 2011; 480: 547–51.

44 Monzio Compagnoni G, Kleiner G, Samarani M, Aureli M, Faustini G, Bellucci A et al. Mitochondrial Dysregulation and Impaired Autophagy in iPSC-Derived Dopaminergic Neurons of Multiple System Atrophy. Stem Cell Reports 2018; 11: 1185–1198.

